# Measures of time series coupling based on generalized weighted multiple regression

**DOI:** 10.1101/235721

**Authors:** Roberto D. Pascual-Marqui, Rolando J. Biscay, Jorge Bosch-Bayard, Pascal Faber, Toshihiko Kinoshita, Kieko Kochi, Patricia Milz, Keiichiro Nishida, Masafumi Yoshimura

## Abstract

The sharing and the transmission of information between cortical brain regions is carried out by mechanisms that are still not fully understood. A deeper understanding should shed light on how consciousness and cognition are implemented in the brain. Research activity in this field has recently been focusing on the discovery of non-conventional coupling mechanisms, such as all forms of cross-frequency couplings between diverse combinations of amplitudes and phases, applied to measured or estimated cortical signals of electric neuronal activity. However, all coupling measures that involve phase computation have poor statistical properties. In this work, the conventional estimators for the well-known phase-phase (phase synchronization or locking), phase-amplitude, and phaseamplitude-amplitude couplings are generalized by means of the weighted multiple regression model. The choice of appropriate weights produces estimators that bypass the need for computing the complex-valued phase. In addition, a new coupling, denoted as the inhibitory coupling (InhCo), is introduced and defined as the dependence of one complex-valued variable on the inverse and on the conjugate inverse of another complex-valued variable. A weighted version denoted as wInhCo is also introduced, bypassing the need for computing the inverse of a complex variable, which has very poor statistical properties. The importance of this form of inhibitory coupling is that it may capture well-known processes, such as the observed inverse alpha/gamma relation within the same cortical region, or the inverse alpha/alpha relation between distant cortical regions.

## 2. Introduction

The sharing and the transmission of information between cortical brain regions is carried out by mechanisms that are still not fully understood. This problem is of great interest, because it may shed light on how consciousness and cognition are implemented in the brain. Research activity in this field has recently been focusing on the discovery of non-conventional coupling mechanisms, such as all forms of cross-frequency couplings between diverse combinations of amplitudes and phases, applied to measured or estimated cortical signals of electric neuronal activity. For a review, see e.g. Jirsa and Muller (2013).

In this work, the conventional estimators for the well-known phase-phase (phase synchronization or locking), phase-amplitude, and phase-amplitude-amplitude couplings are generalized by means of the weighted multiple regression model. The choice of appropriate weights produces estimators that bypass the need for computing the complex-valued phase, which has very poor statistical properties.

In addition, a new coupling, denoted as the inhibitory coupling (InhCo), is introduced and defined as the dependence of one complex-valued variable on the inverse and on the conjugate inverse of another complex-valued variable. A weighted version denoted as wInhCo is also introduced, bypassing the need for computing the inverse of a complex variable, which has very poor statistical properties. The importance of this form of inhibitory coupling is that it may capture well-known processes, such as the observed inverse alpha/gamma relation within the same cortical region, or the inverse alpha/alpha relation between distant cortical regions (see e.g. De Pesters et al 2016).

## 3. Complex-valued random variables

Let *x_i_,y_i_* ∈ℂ, for *i* = 1…*N*, denote a paired sample of size *N,* of complex-valued random variables, centered to zero-mean.

Two examples for this type of data are:

1. The collection of Fourier coefficients at discrete frequencies (ω_*x*_, ω_*y*_), for two stationary signals (*x*, *y*), for which *N* epochs are available.
2. The complex-valued analytic signals obtained with the Hilbert transform at frequency bands (*b_x_*, *b_y_*), for two signals (*x*, *y*), for *N* time samples.

A classical textbook on the statistics of stationary processes in the frequency domain can be found e.g. in Brillinger 2001.

A paper on how to compute in practice the discrete Hilbert transform (rather than its actual definition as the principal value of an integral) and the analytic signal can be found in e.g. Marple 1999.

Given complex random variables *z_i_*, for *i* = 1…*N*, centered to zero-mean, the non-negative real-valued amplitude is denoted as:

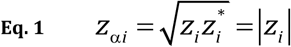

Its complex-valued phase is denoted as:

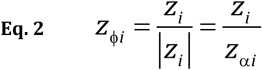

Its inverse can be expressed as:

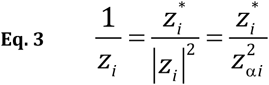

And the inverse of the conjugate variable can be expressed as:

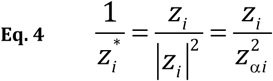

In general, the superscript ^“*”^ denotes complex conjugate and vector-matrix transposed; and |•|denotes the norm of the argument.

The centered, zero-mean amplitude is denoted:

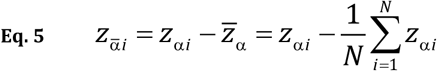

## 4. The linear model and the coherence

Consider the linear model for the zero-mean complex-valued random variables (*x*,*y*):

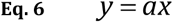

The simplest measure of coupling in this case is the classical complex-valued coherence (see e.g. Brillinger 2001):

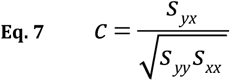

with:

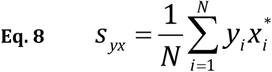

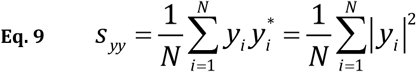

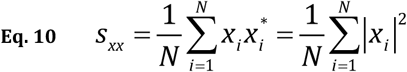

## 5. The widely linear model and the squared multiple coherence

The widely linear model for the zero-mean complex-valued random variables (*x*, *y*) is:

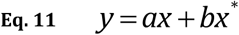

which was proposed by Picinbono and Chevalier (1995). This type of model (Eq. 11) has been extensively studied in, see e.g. Mandic and Goh (2009), and Schreier and Scharf (2010).

A measure of coupling for this case is the squared multiple coherence of the response *“y”* with the predictors “*x*” and “*x*^*^”.

In general, the squared multiple coherence (correlation) for centered, zero-mean variables, with response ϑ ∈ ℂ (ϑ ∈ ℝ) and *“p”* predictors **Ψ** ∈ ℂ^*p*×1^ (**Ψ** ∈ ℝ^*p*×1^) is:

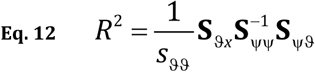

where in general the covariance matrix is:

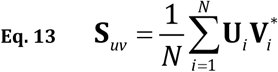

for any two zero-mean variables **U** ∈ ℂ^*q*×1^) and **V** ∈ ℂ^*r*×1^ (**V** ∈ ℝ^*r*×1^.

The multiple squared coherence for the widely linear model in Eq. 11 is calculated for the variables:

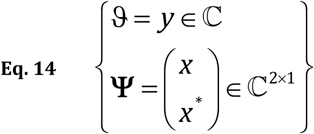

## 6. Models based on relations between different combinations of phases and amplitudes of complex variables

### 6.1. Phase-phase coupling (a.k.a. phase synchronization and phase locking value)

Two definitions for phase-phase coupling will be considered.

In a first case, it is defined as the average complex valued phase difference:

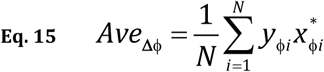

or its modulus |*Ave*_Δϕ_| or squared modulus |*Ave*_Δϕ_|^2^, which lie in the range zero to one. A value of one is attained when the phase difference is constant, and a value of zero can be attained when the phase difference is uniformly distributed. This definition corresponds to the concept of phase-coupling as proposed by Rosenblum et al (1996).

In a second case, it can be modeled as a linear regression between phase variables, for which abundant theory exists, see e.g. Jupp and Mardia (1980) and Mardia and Jupp (2000). For such a linear relation between the two phase variables, the coherence is:

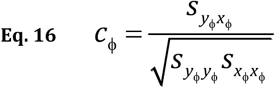

Ultimately, both definitions lead to the same final result, with:

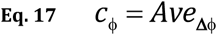

However, these two definitions lead to different weighted estimators, as will be shown below.

### 6.2. Phase-amplitude coupling (PAC)

Phase-amplitude coupling (PAC) refers to the dependence of the real-valued amplitude of one variable on the complex-valued phase of another variable (which can correspond to two different time series or to the same time series). Many different PAC measures have been proposed in the literature, as reviewed by, e.g. Penny et al (2008) and van Wijk et al 2015.

One PAC model is:

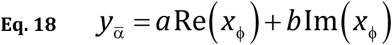

where 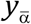 denotes the centered zero-mean amplitude, Re(•) and Im (•) denote the real and imaginary parts of the argument, and (*a*, *b*) are the real-valued regression coefficients. This general linear model was proposed in Penny et al (2008). For this model, PAC can be quantified by the squared multiple correlation coefficient (Eq. 12) for the variables:

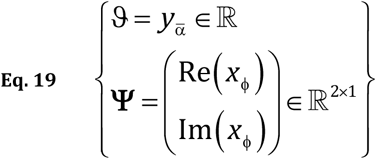

The multiple squared correlation (Eq. 12) for this PAC model (Eq. 19) will be denoted as 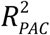.

A generalization of the PAC model, denoted here as the PAAC model, was given in van Wijk et al 2015, which considers the dependence of the real-valued amplitude of one variable on the complex-valued phase and the real-valued amplitude of another variable:

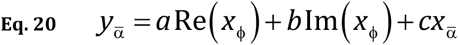

where 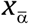 denotes the centered, zero-mean amplitude, and (*a,b,c*) are the real-valued regression coefficients. In this case (PAAC) there are three regression coefficients, with the coefficients (*a*, *b*) corresponding to the “phase-amplitude coupling” component, and with the coefficient (*c*) corresponding to the “amplitude-amplitude coupling” component. Measures of coupling can be defined by the regression coefficients (*a,b,c*), and in general by the squared multiple correlation coefficient for the response 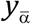 as a function of the predictors Re(*x*_ϕ_), Im(*x*_ϕ_), and 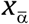, i.e. using (Eq. 12) for the variables:

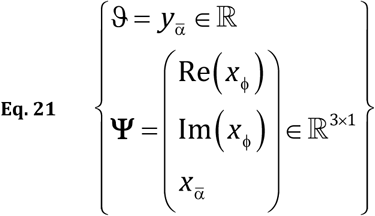

The multiple squared correlation (Eq. 12) for this more general PAC model (Eq. 21) will be denoted as 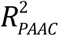.

### 6.3. Inhibitory coupling (InhCo)

We define “inhibitory coupling” (InhCo) as the dependence of one complex-valued variable on the inverse and on the conjugate inverse of another complex-valued variable:

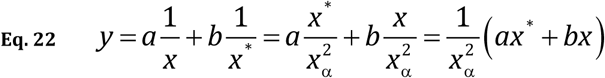

For zero-mean variables (*x*,*y*), the squared multiple coherence for the response variable “*y*” as a function of the predictors 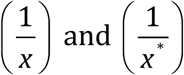 quantifies the InhCo, with:

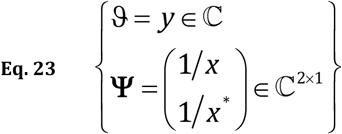

for Eq. 12.

The use of the term “inhibitory” corresponds to the notion of one process inhibiting another, such as the observed inverse alpha/gamma relation within the same cortical region, or the inverse alpha/alpha relation between distant cortical regions (see e.g. De Pesters et al 2016).

## 7. Instability of complex-valued phase and inverse variables

The computation of a complex-valued phase variable, as defined in Eq. 2, requires a non-zero amplitude value. In practice, when the amplitude is near zero, the phase is nearly undefined. This affects the estimated phase-phase coupling (Eq. 15, Eq. 16), and all forms of phase-amplitude coupling (see e.g. Eq. 18 and Eq. 20). In a similar manner, the computation of the inverse of a complex-valued variable requires a non-zero amplitude value, and is nearly undefined for near-zero amplitudes, which would affect the estimated inhibitory coupling (InhCo) in Eq. 22.

Little attention has been given to this problem in the literature, as reviewed by Kovach (2017). In Kovach (2017), the effect of low amplitudes on phase-phase coupling was studied in detail, and a solution was proposed in the form of an amplitude weighted version of the phase-phase coupling. The Kovach (2017) estimator is presented in a later section of this present study.

## 8. General stabilization by setting a threshold for the minimum allowed amplitude

One naïve solution that immediately comes to mind is to set a threshold for the minimum allowed amplitude, and to discard all data below the threshold, prior to estimating the coupling measures.

Note that for stationary time series, the squared amplitude of the Fourier transform is the periodogram, which has an asymptotic chi-square distribution with two degrees of freedom, of the 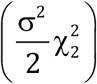, form where σ^2^ is the population spectral density. Standardized squared amplitudes, defined as the squared amplitudes divided by the average, have an approximate 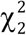 distribution.

In this case, the p=0.05 threshold occurs at a value of 0.103. Thus, 0.103 can be taken as an approximate minimum allowed standardized squared amplitude value, such that any standardized squared amplitude below this threshold should be discarded and not included in the estimation of coupling models that involve phase or inverse variables.

For instance, an algorithm for computing phase-phase coupling is:

Step#1: Given the original data (*x_i_, y_i_*) for *i* = 1…*N*. And given a threshold for the minimum allowed squared amplitude denoted as min*Amp*^2^ (e.g. min*Amp*^2^ = 0.103).
Step#2: Compute the squared amplitudes 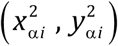.
Step#3: Compute the mean values for the squared amplitudes, denoted as 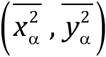.
Step#4: Transform the squared amplitudes to standardized squared amplitudes 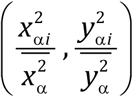.
Step#5: For *i* = 1…*N* do if 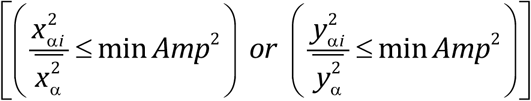 then delete the i-th pair (*x_i_, y_i_*).
Step#6: With a possibly reduced data set, then compute the phase-phase coupling (Eq. 15, Eq. 16).

The algorithm for PAC and for InhCo (Eq. 18, Eq. 20, Eq. 22) is basically the same:

Step#1’: Given the original data (*x_i_, y_i_*) for *i* = 1…*N*. And given a threshold for the minimum allowed squared amplitude denoted as **min** *Amp*^2^ (e.g. **min** *Amp*^2^ = **0.103**).
Step#2’: Compute the squared amplitudes 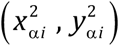.
Step#3’: Compute the mean values for the squared amplitudes for 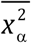.
Step#4’: Transform the squared amplitudes to standardized squared amplitudes 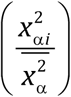.
Step#5’: For *i* = 1…*N* do if 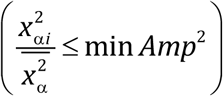 then delete the i-th pair (*x_i_, y_i_*).
Step#6’: With a possibly reduced data set, then compute PAC and InhCo (Eq. 18, Eq. 20, Eq. 22).

## 9. Phase-phase coupling stabilization of by the method of weighted averages (Kovach 2017)

The phase-phase coupling in Eq. 15 has the form of an average:

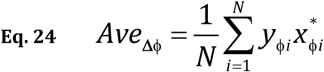

To avoid the effect of low amplitudes when computing phases, Kovach (2017) proposed a weighted average:

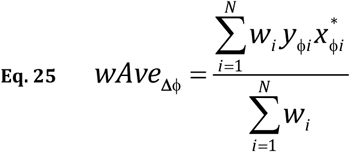

with non-negative weights:

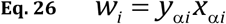

which gives the amplitude-weighted phase locking value of Kovach (2017):

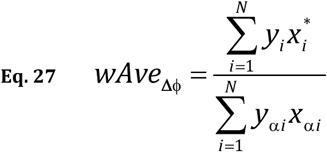

The advantage of the weighted average in Eq. 27 is that it does not require the computation of phases. In addition, data with low amplitude contributes less to the coupling measure.

However, this stabilization method cannot be generalized to other coupling measures, such as those used for PAC and InhCo couplings, which have the form of a squared multiple correlation or a squared multiple coherence. This is because these measures cannot be explicitly expressed as a simple average that can then be conveniently weighted.

Nevertheless, the basic idea of finding appropriate weights for a weighted average as proposed by Kovach (2017), can be extended to develop new estimators in the form of weighted correlations and coherences with appropriate weights. The new estimators follow next.

## 10. General stabilization by weight transformations

Consider centered, zero-mean random variables, with response υ ∈ ℂ (υ ∈ ℝ) and *“p”* predictors **ϒ** ∈ ℝ^p×1^, with sample data (υ_*i*_, **ϒ**_*i*_, for *i* = 1…*N*. And let *w_t_* ≥0 denote real-valued, nonnegative weights, such that:

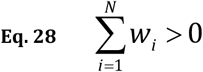

The weighted multiple coherence (correlation) can be written as:

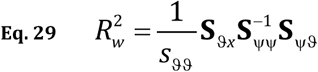

where:

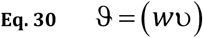

and

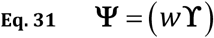

where the covariance matrices are defined in general in Eq. 13.

Note that Eq. 29 is invariant to a change of global scale of the weights, i.e. the weighted multiple coherence (correlation) does not change for new weights 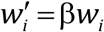, for any constant β> 0.

### 10.1. Weighted phase-phase coherence

Using the notation for the weighted multiple regression model corresponding to Eq. 29, Eq. 30, and Eq. 31, the complex-valued phase variables are:

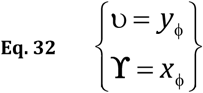

The choice of weights in this case is the same as used by Kovach (2017) in Eq. 26:

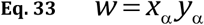

This gives:

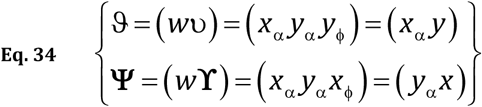

which are to be used in Eq. 29. Because this is a simple bivariate case, the weighted phase-phase squared coherence and coherence can be explicitly expressed as:

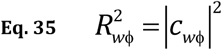

with:

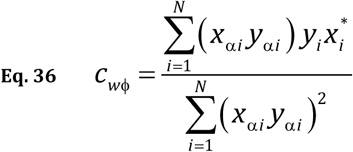

Note that the weighted phase-phase coherence introduced in this work as Eq. 36, differs from weighted average of Kovach (2017) in Eq. 27, in that the coherence approach in Eq. 36 gives more importance to phase differences that have higher joint amplitudes.

However, both approaches have the main common feature of avoiding the instability related to computing complex-valued phases.

### 10.2. Weighted phase-amplitude coupling (wPAC)

Using the notation for the weighted multiple regression model corresponding to Eq. 29, Eq. 30, and Eq. 31, the PAC model in Eq. 18 has real variables:

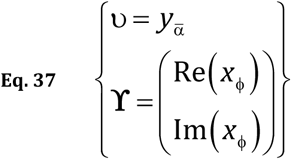

The obvious stabilizing choice of weights in this is:

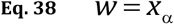

This gives:

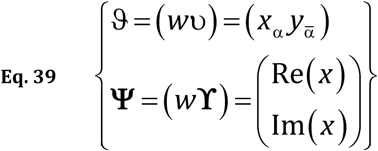

which are to be used in Eq. 29, giving:

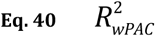

which is the weighted squared multiple correlation for the PAC model.

This does not require the computation of phases, and does not have the statistical instability of the non-weighted model. In explicit form, the weighted model is:

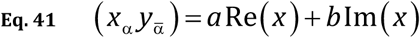

### 10.3. Weighted phase-amplitude-amplitude coupling (wPAAC)

Using the notation for the weighted multiple regression model corresponding to Eq. 29, Eq. 30, and Eq. 31, the PAAC model in Eq. 20 has real variables:

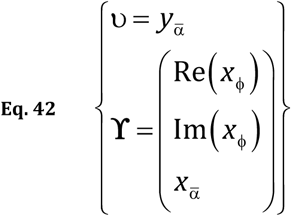

The obvious stabilizing choice of weights in this is the same as before, see Eq. 38. This gives:

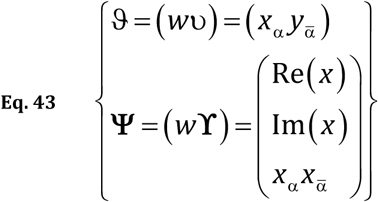

which are to be used in Eq. 29, giving:

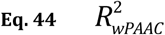

which is the weighted squared multiple correlation for the PAAC model.

This does not require the computation of phases, and does not have the statistical instability of the non-weighted model. In explicit form, the weighted model is:

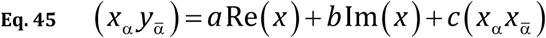

### 10.4. Weighted inhibitory coupling (wInhCo)

Using the notation for the weighted multiple regression model corresponding to Eq. 29, Eq. 30, and Eq. 31, the InhCo model in Eq. 22 has complex variables:

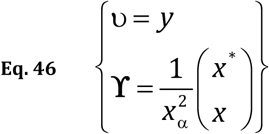

The obvious stabilizing choice of weights in this is:

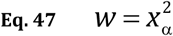

This gives:

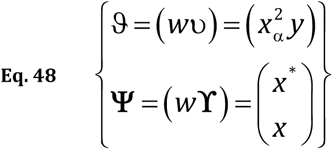

which are to be used in Eq. 29, giving:

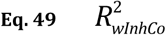

which is the weighted squared multiple coherence for the InhCo model.

This does not require the computation of inverse of complex variables, which would produce statistical instability in the non-weighted model. In explicit form, the weighted model is:

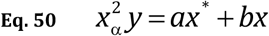

## 11. Conclusions

Coupling measures that involved complex phase or complex inverse variables have very poor statistical properties, This was shown by Kovach (2017) for phase-phase coupling. One remedy, as proposed by Kovach (2017), was to replace the average with a weighted version, that bypassed the need of computing the complex phase.

Motivated by this methodology, but applying it to weighted multiple regression models, we present weighted versions of many forms of couplings between diverse combinations of phases and amplitudes and inverses. Based on the same arguments as in Kovach (2017), these new weighted estimators should have improved statistical properties as compared with the estimators commonly used in the literature up to now.

## References

Brillinger, D.R., 2001. Time series: data analysis and theory. Society for Industrial and Applied Mathematics.

Jirsa, V., and Muller, V. (2013). Cross-frequency coupling in real and virtual brain networks. Front Comput Neurosci 7, 78.

Jupp, P.E. and Mardia, K.V., 1980. A general correlation coefficient for directional data and related regression problems. Biometrika, 67(1), pp.163–173.

Kovach, C., 2017. A biased look at phase locking: Brief critical review and proposed remedy. IEEE Transactions on Signal Processing, 65 (17), pp. 4468–4480.

Mandic, D.P. and Goh, V.S.L., 2009. Complex-valued nonlinear adaptive filters: noncircularity, widely linear and neural models (Vol. 59). John Wiley & Sons.

Mardia, K.V. and Jupp, P.E., 2000. Directional statistics (Vol. 494). John Wiley & Sons.

Marple, L., 1999. Computing the discrete-time” analytic” signal via FFT. IEEE Transactions on signal processing, 47(9), pp.2600–2603.

Penny WD, Duzel E, Miller KJ, Ojemann JG. Testing for nested oscillation. Journal of neuroscience methods. 2008 Sep 15;174(1):50–61.

De Pesters, A., Coon, W.G., Brunner, P., Gunduz, A., Ritaccio, A.L., Brunet, N.M., De Weerd, P., Roberts, M.J., Oostenveld, R., Fries, P., and Schalk, G. (2016). Alpha power indexes task-related networks on large and small scales: A multimodal ECoG study in humans and a non-human primate. Neuroimage.

Picinbono, B. and Chevalier, P., 1995. Widely linear estimation with complex data. IEEE transactions on Signal Processing, 43(8), pp.2030–2033.

Rosenblum MG, Pikovsky AS, Kurths J. Phase synchronization of chaotic oscillators. Physical review letters. 1996 Mar 11;76(11):1804.

Schreier, P.J. and Scharf, L.L., 2010. Statistical signal processing of complex-valued data: the theory of improper and noncircular signals. Cambridge University Press.

van Wijk BC, Jha A, Penny W, Litvak V. Parametric estimation of cross-frequency coupling. Journal of neuroscience methods. 2015 Mar 30;243:94–102.

